# A functional spiking-neuron model of activity-silent working memory in humans based on calcium-mediated short-term synaptic plasticity

**DOI:** 10.1101/823559

**Authors:** Matthijs Pals, Terrence C. Stewart, Elkan G. Akyürek, Jelmer P. Borst

## Abstract

In this paper, we present a functional spiking-neuron model of human working memory (WM). This model combines neural firing for encoding of information with activity-silent maintenance. While it used to be widely assumed that information in WM is maintained through persistent recurrent activity, recent studies have shown that information can be maintained without persistent firing; instead, information can be stored in activity-silent states. A candidate mechanism underlying this type of storage is short-term synaptic plasticity (STSP), by which the strength of connections between neurons rapidly changes to encode new information. To demonstrate that STSP can lead to functional behavior, we integrated STSP by means of calcium-mediated synaptic facilitation in a large-scale spiking-neuron model and added a decision mechanism. The model was used to simulate a recent study that measured behavior and EEG activity of participants in three delayed-response tasks. In these tasks, one or two visual gratings had to be maintained in WM, and compared to subsequent probes. The original study demonstrated that WM contents and its priority status could be decoded from neural activity elicited by a task-irrelevant stimulus displayed during the activity-silent maintenance period. In support of our model, we show that it can perform these tasks, and that both its behavior as well as its neural representations are in agreement with the human data. We conclude that information in WM can be effectively maintained in activity-silent states by means of calcium-mediated STSP.

**Author Summary:** Mentally maintaining information for short periods of time in working memory is crucial for human adaptive behavior. It was recently shown that the human brain does not only store information through neural firing – as was widely believed – but also maintains information in activity-silent states. Here, we present a detailed neural model of how this could happen in our brain through short-term synaptic plasticity: rapidly adapting the connection strengths between neurons in response to incoming information. By reactivating the adapted network, the stored information can be read out later. We show that our model can perform three working memory tasks as accurately as human participants can, while using similar mental representations. We conclude that our model is a plausible and effective neural implementation of human working memory.

## Introduction

The ability to temporarily hold information in working memory (WM) is a crucial part of day-to-day life: it is what allows us to remember someone’s name at a cocktail party, what ingredients to buy at the supermarket for dinner, and which platform we need to go to when changing trains [1,2]. The maintenance of information in WM is often studied by means of a delayed-response task, in which a briefly presented memory item is followed by a delay period [3,4]. The delay period ends with the presentation of a probe that the participants need to compare to the memorized item. The maintenance of information during the delay period of such tasks was long thought to be mediated by continuously spiking neurons [5,6]. Although neural spiking is certainly important for WM, it was recently shown that spiking activity during delay periods can be intermittent or even absent [7–11]. This suggests that information may be stored instead using activity-silent mechanisms, for instance through transient connectivity patterns in the brain [2,12,13]. The spiking activity observed previously might reflect the initial phase necessary to initialize new synaptic weights, active maintenance of the focus of attention [14–17], or the read-out of information from working memory [13,18].

One of the candidate mechanisms for storing information in activity-silent states is short-term synaptic plasticity [STSP; 19], which entails rapid changes in the strength of connections between neurons to reflect new information being presented to the network [12]. Indeed, it was previously shown that synapses in areas implicated in WM can be (temporarily) strengthened (or *facilitated*; [20,21]), potentially as a consequence of residual calcium building up in presynaptic terminals [19,22]. In this way residual calcium effectively leaves a ‘synaptic trace’ of what is currently stored in WM. An elegant implementation of activity-silent storage by means of STSP was proposed by Mongillo and colleagues [12], who developed a model that can maintain information through calcium-mediated synaptic facilitation in recurrent networks of simulated spiking neurons. In response to a particular input to the network, a subset of the neurons fires, with the result that their outgoing connections are facilitated. Subsequently, stored information can be read out by applying a network-wide non-specific input that will be mostly subthreshold for non-facilitated neurons but leads to firing of facilitated neurons.

In the current study, we show that the mechanism proposed by Mongillo and colleagues [12] not only results in efficient and robust storage, but also in effective, functional human behavior. Although previous models have suggested that this should be the case, these models were based on rate neurons and did not attempt to match human behavior [8,18]. Here, we integrated the calcium-mediated STSP mechanism in a large-scale spiking-neuron model that can perform a delayed-response task. To have the model make the necessary decisions, we introduced a mechanism that effectively compares newly presented information to what is currently held in WM, inspired by Myers’ template matching proposal [23]. To evaluate this model, we used three previously reported datasets of visual WM tasks, in two of which activity-silent memory states were measured by means of electroencephalography (EEG) [3]. To this end, Wolff and colleagues developed an innovative method to probe activity-silent brain states [3,24]. They showed that when the WM network is perturbed by a high-contrast task-neutral stimulus during maintenance, ensuing neural activity reveals what is currently held in an activity-silent state.

In their main experiment (Experiment 1; [3]), each trial started with the display of two randomly oriented gratings (Fig 1). After an 800 ms fixation period, this was followed by a cue indicating which of the two stimuli had to be maintained in memory. In order to examine the contents of WM during the subsequent delay part of the trial, an impulse stimulus was presented 900 ms later. At the end of each trial, participants had to indicate whether a probe stimulus was rotated clockwise or counter-clockwise with respect to the cued memory item. To track the contents of WM, a decoding analysis was applied to the EEG data [3]. It was shown that decoding accuracy quickly dropped to chance level after presentation of the memory items, but returned when the probe was presented. This indicates that between the presentation of the memory items and the probe, information is maintained in an activity-silent (or at least quiescent) state. In addition, it was shown that it is possible to decode the orientation of the cued memory item from the EEG data during maintenance in response to the impulse stimulus. Thus, when the WM network was perturbed by a task-neutral stimulus, the ensuing signal allowed for decoding of the current contents of the activity-silent state. Interestingly, at this point in the trial, only the orientation of the cued memory item could be decoded, indicating that the uncued stimulus was quickly forgotten, or actively cleared from memory.

**Fig 1.**
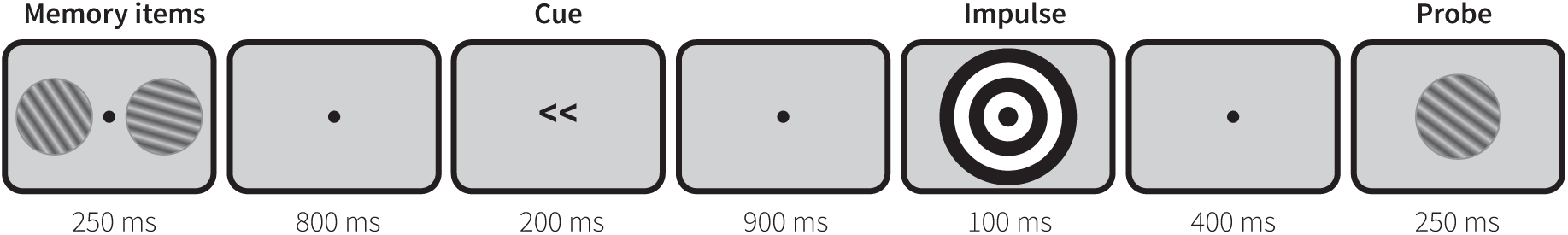
The retro-cue delayed-response task. After the presentation of the memory items, a cue indicates which grating needs to be maintained in WM for judgement of the probe. Decodable EEG activity is elicited by the task-neutral impulse, which is presented between the cue and the probe.

To evaluate our model, we let it perform the same experiment – including the application of the impulse perturbation method – and compared both our model’s performance as well as its mental representations and underlying spiking behavior to the human data. To further characterize the model, we then performed a parameter exploration and tested whether the model generalizes to other experimental setups. For the latter, we applied it to Experiments 2 and 3 of the Wolff study [3], in which they investigated the maintenance of multiple items in WM and the potential disruptive effect of longer impulse stimuli, respectively.

## Results

### Model Architecture

In order to implement a functional spiking-neuron model of WM we used Nengo, a framework for building large-scale brain models that link single cell activity to demonstrative cognitive abilities [25–27]. In this framework, information is represented by vectors of real numbers that can be encoded and decoded from the collective spiking activity of a population of neurons. Connections between neural populations allow for both communication and transformation of information. Here, Nengo acts as a ‘neural compiler’: given a desired function, the connection weight matrix between populations is calculated so that this function is approximated. Besides pre-calculating connection weights, plasticity can be introduced by making use of biologically plausible learning rules [28].

To account for short-term synaptic plasticity, we integrated the calcium kinetics mechanism proposed by Mongillo and colleagues [12] in the model. Accordingly, synaptic efficiency between two neurons is dependent on two parameters: the amount of available resources to the presynaptic neuron (reflecting neurotransmitters) and the presynaptic calcium level. Each time a neuron fires, the amount of available resources decreases, reducing synaptic efficiency. As resources are quickly replenished (in the order of 200 ms), this results in short-term depression of firing rates. However, at the same time calcium flows into the presynaptic terminals, *increasing* synaptic efficiency. Because calcium is much slower to return to its baseline levels than the resources, the synaptic connection is facilitated in the long-term, for about 1.5 seconds (see Methods for the effects of parameter settings). To calculate the final synaptic strength of each connection, we multiplied the momentary synaptic efficiency with precalculated connection weights (Equation 1.3).

This STSP mechanism was applied to the recurrent connections of two working memory populations in our model. As described above, the aim is to simulate a delayed-response task in which the orientation of two memory items has to be compared to a probe ([3]; Fig 1). In this task, significant EEG lateralization was observed at posterior electrodes after presentation of the cue. We therefore hypothesized that distinct populations of neurons are responsible for processing visual stimuli presented in the left and right visual field. Correspondingly, the model was divided into two independent modules, each responsible for perceiving and representing one of the two incoming stimuli (Fig. 2).

**Fig 2.**
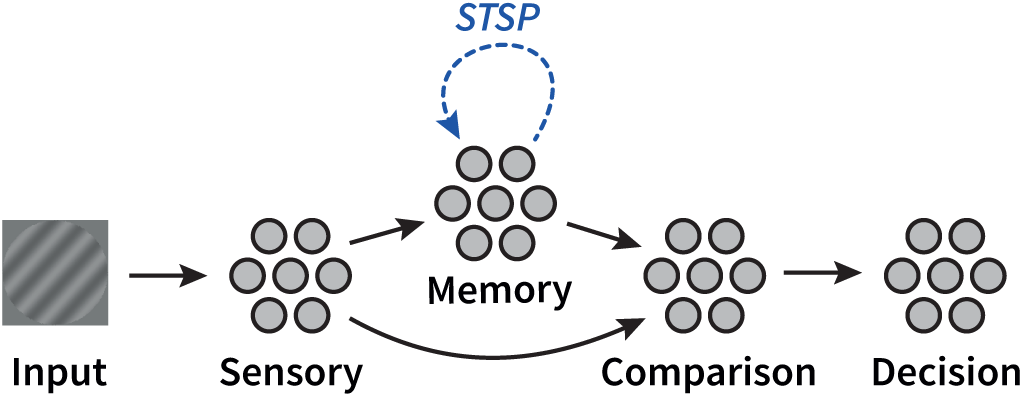
Model architecture. The model is divided in two modules (only one is pictured) representing the two visual hemispheres. Stimuli enter via a sensory population that transforms the input into a vector. This vector is then sent to a recurrently connected memory population exhibiting STSP. The comparison population integrates information from both the sensory and memory ensembles, the result of which is interpreted by the decision population.

In order to demonstrate that our model is able to deal with real-world input, the actual stimuli from [3] were presented to the model. The sensory populations use two-dimensional Gabor filters as encoders [26,29,30]. As a result, the information present in the gratings – including their direction – is encoded into 24-dimensional vectors that are passed on to the memory populations. That is, the information encoded into the neurons is a compressed representation of the input image, using the top 24 singular values as per SVD (see Methods for more details). The memory populations contain recurrent connections exhibiting STSP, in line with previous models of WM and anatomical areas associated with WM [e.g., 21,31,32]. Consequently, the first stimulus during a trial will drive facilitation of recurrent connections representing this stimulus. Neural activity resulting from subsequent stimuli will be affected by this change in connectivity.

This enables implementing decision making as a match-filter process [8,13]. To decide on the orientation change of the probe compared to the relevant memory item, both the sensory and memory populations communicate the orientation of the gratings to a comparison population. When a probe is presented, the orientation received from the sensory population is driven entirely by the incoming stimulus. However, the orientation of the memory population is driven by a dynamic combination of activity resulting from the incoming stimulus and activity from facilitated connections as a result of the encoded memory item (i.e., hysteresis). In other words, the orientation represented by the memory population reflects the orientation of the probe ‘tuned’ by the orientation of the memory item stored in facilitated synapses, over time reverting to the new probe stimulus. To estimate the orientation difference between the memory item and the probe, the outgoing connections from the comparison layer subtract the two represented orientations. The resulting one-dimensional value reflects the signed difference between the orientation of the memory item and that of the probe stimulus.

### Neural representations

The model simulated the experiment reported in [3] and illustrated in Fig 1. In the original paper, it was shown that decoding accuracy quickly dropped after presentation of the memory items but returned during presentation of the probe – as would be expected for an activity-silent maintenance mechanism. Correspondingly, we examined the spiking activity and quality of representations of our model during the task, in order to validate that any maintenance of information in our model is realized in activity-silent states and not by persistent firing.

Fig 3 shows the spiking activity of the neurons in the memory populations of both modules during one trial (A: cued module, B: uncued module), together with the mean amount of resources (*x*) and calcium (*u*) in these populations. In both modules, there is spiking activity during and shortly after presentation of the memory items, the impulse stimulus, and the probe, but not in between. The spiking activity causes the amount of available resources and the calcium level to decrease and increase, respectively. The resulting short-term depression can directly be observed as the amount of spiking declines after the onset of a stimulus, although it periodically reactivates.

**Fig 3.**
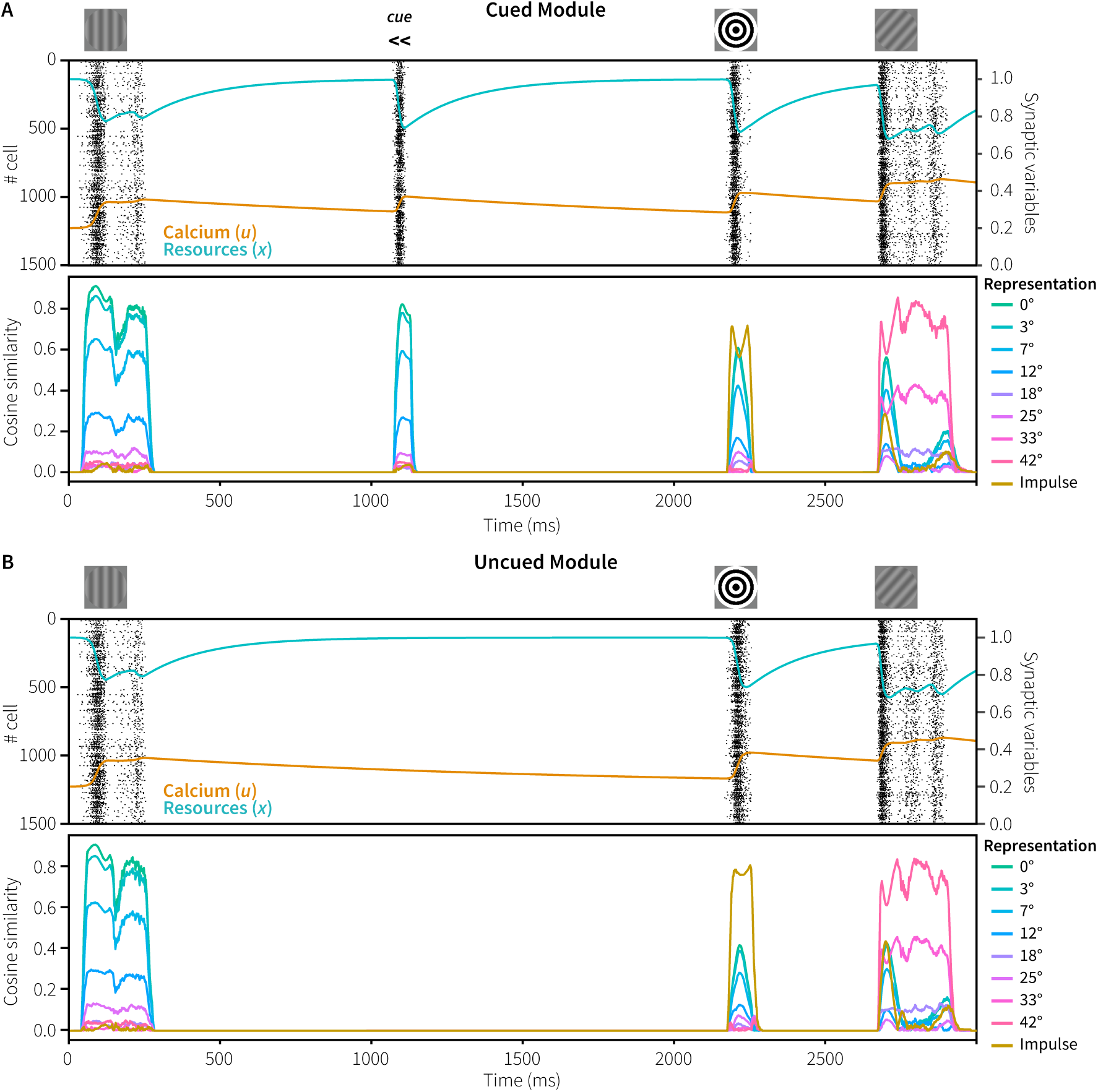
Spiking behavior and representations in Experiment 1. Top: spiking activity of the memory populations of the cued (A) and uncued (B) modules, including resource and calcium levels during a trial. Bottom: absolute normalized cosine similarity between the vector represented by the memory populations and ideal vectors, averaged over 100 trials with 0° memory items and 42° probes with constant within-trial phase.

In the original experiment, a retro-cue that indicated which of the two previously presented items needed to be memorized was presented 800 ms later, which was followed by significant lateralization at posterior electrodes. To mimic this, the memory population of the cued module is briefly reactivated by means of a non-specific population wide input [cf. 12]. This not only re-activates the memory item, but also helps to maintain the stimulus for a longer time period, as reactivation of facilitated synapses will lead to re-facilitation of those connections.

Next, we analyzed the vectors represented by the memory populations of both the cued and uncued module. Fig 3 (bottom of each panel) shows the absolute cosine similarity between the vector represented by the memory populations and the ideal vectors of potential representations. To clearly illustrate the difference between the two modules, the mean cosine similarity was calculated over 100 trials in which both modules were presented with the same memory item and probe, with a rotation of 0° and 42°, respectively. Note that in the simulation of the real experiment, the cued and uncued modules are never presented with the same memory item.

During presentation of the initial memory item of 0°, the vectors represented by both modules are very similar to the ideal 0° vector. In addition, the cosine similarity is inversely correlated with the angular difference between the represented vector of 0° and potential representations, indicating that similar stimuli are represented by similar vectors and firing patterns. As was the case in the original experiment, during the delay periods we could not decode what is being represented by the neural populations as there is no spiking activity – indicating activity-silent memory. However, in response to the non-specific reactivation of the cued model at 1050 ms, there was spiking activity that clearly represents the originally encoded vector. It therefore appears that neural connections representing the memory item were indeed facilitated, and that mainly those connections and neurons get activated in response to the non-specific reactivation elicited by the cue.

One of the main results of the original study was that the EEG activity in response to the impulse stimulus contained the orientation of the cued memory item, and not of the uncued item [3]. This was taken to show that a stimulus is only maintained in an activity-silent state if it is still needed for the task. If not, it is quickly forgotten or actively cleared from the network. To see if our model has both the same storage and forgetting capabilities, we examined the vectors represented by the memory populations of the cued and uncued module during presentation of the impulse (Fig 4; cf. Fig 3, 2150-2300 ms). In both modules, the memory populations start representing the impulse stimulus. When the facilitated recurrent connections of the cued and uncued items become activated, both modules also represent the original 0° memory item. However, only for the cued module does the represented vector become (very briefly) more similar to the ideal memory item than to the impulse vector. In contrast, in the uncued model the represented vector is twice as similar to the impulse than to the original memory item, offering a potential explanation of why only the cued, and not the uncued memory item, could be decoded after the impulse [3]. Note that we cannot compare these results directly to the decoding results of Wolff and colleagues, as it is non-trivial to decode model data meaningfully. However, it does illustrate a potential cause of the reported decoding results.

**Fig 4.**
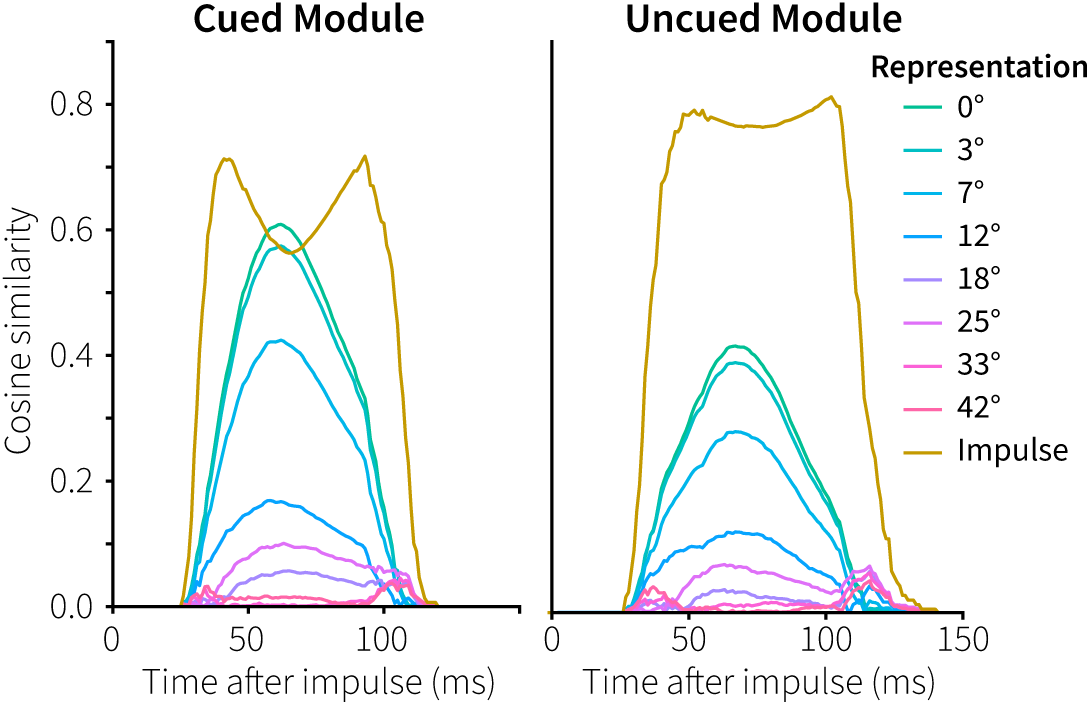
Cued and uncued memory representations in response to the impulse stimulus in Experiment 1. Absolute normalized cosine similarity between the representations in the memory populations and ideal vectors in response. The memory item presented before the impulse had a rotation of 0°.

To summarize: in both the cued and the uncued modules, STSP encodes the initial stimulus. In the cued model, facilitated connections are re-facilitated at the moment of cue, counteracting the gradual calcium decay that goes on in both modules (Fig 3). As a result, once the impulse arrives, only the cued model has sufficiently facilitated connections specific to the memory item to generate a response larger than the impulse representation (Fig 4). Note that the uncued memory population was not actively cleared, but that the calcium levels of the facilitated synapses simply decayed away as it was not reactivated at the moment of the cue.

### Behavior

In order to see if our model not only matches neural activity, but also gives rise to functional behavior similar to human participants, we evaluated its performance. First, to see if the information maintained in the facilitated synapses can be used to produce a relevant response, we inspected the value represented by the decision population in the cued module. This population receives the angular difference between the memory item and the probe from the comparison population, and thus represents a measure of difference between the orientations decoded from the sensory and memory populations. Fig 5 shows the represented value for the possible orientation differences between memory items and probes, averaged over all simulated trials. First, it takes a moment for the probe information to reach this population. Second, the facilitated synapses become activated, reactivating the memory item, thereby leading to different representations in the sensory and memory populations, and thus to a difference in the decision population. Finally, the probe starts overriding the memory representation, reducing the difference until both populations represent the probe and the difference has disappeared. Overall, both the sign and magnitude of the orientation difference are clearly represented in the decision signal.

**Fig 5.**
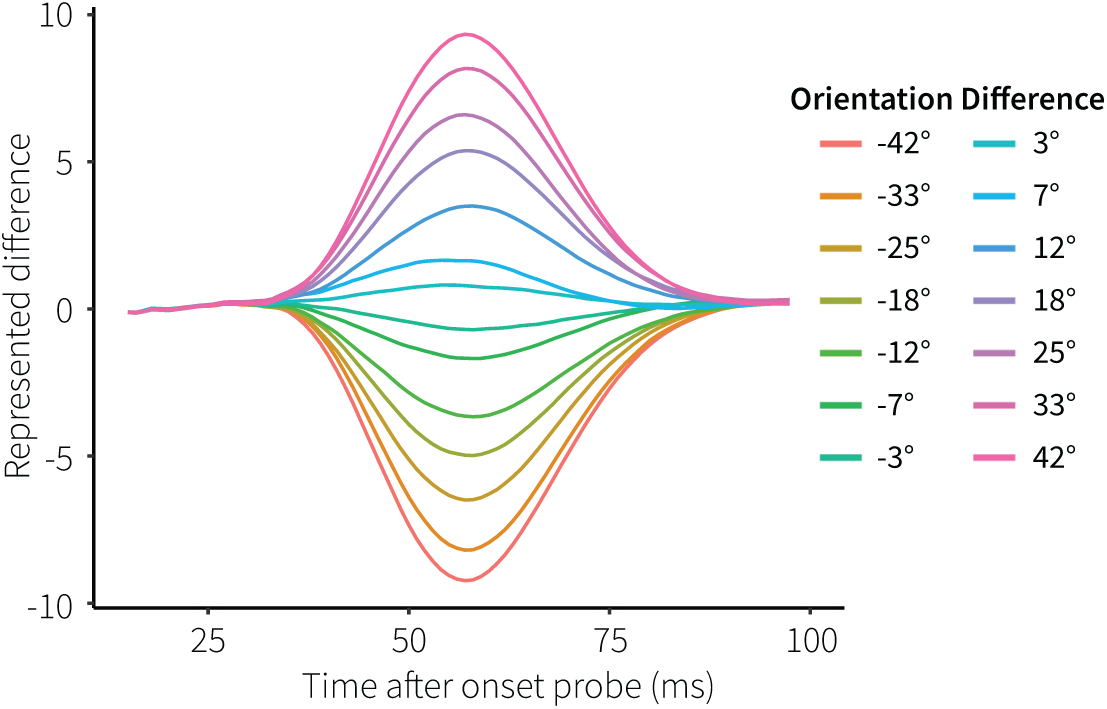
Represented difference in the decision population. Colors indicate experimental orientation differences, averaged over all trials.

To translate this decision signal into a response, we integrated the decision activation after the presentation of the probe. Integrating the evidence corresponding to two distinct decisions has been widely used before in accumulator models of perceptual decision making [e.g., 33]. We did not model motor processes, but simply interpreted a positive result as a clockwise response and a negative result as a counter-clockwise response. Fig 6 shows that the model’s proportion of clockwise responses across orientation differences follows a similar S-shape as the human responses.

**Fig 6.**
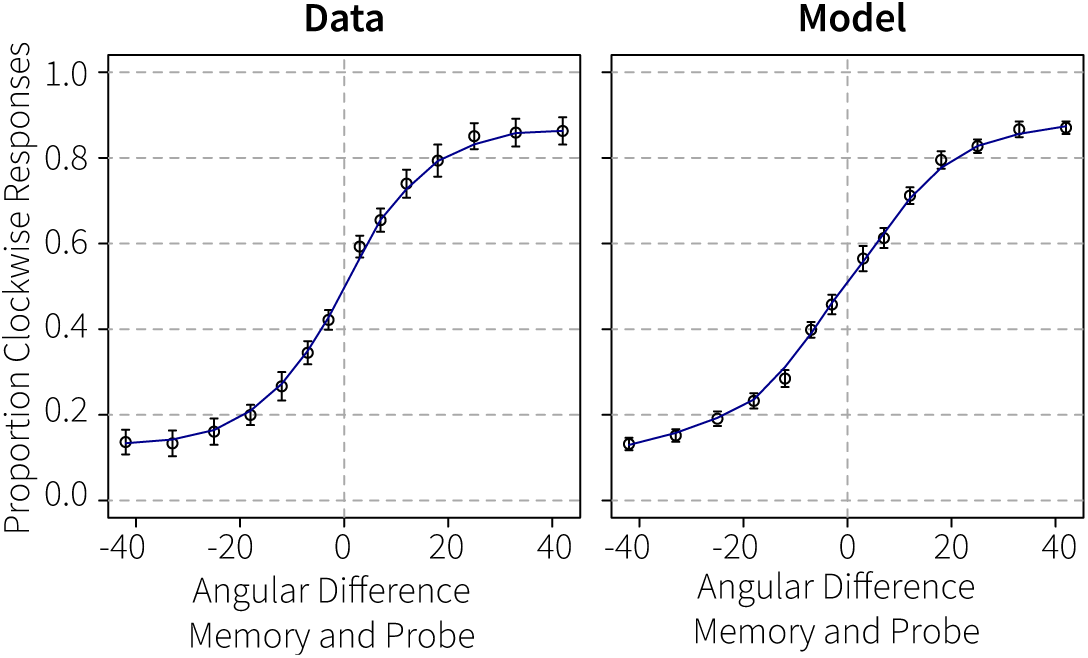
Performance in Experiment 1. Proportion of clockwise responses dependent on the angular difference between the cued memory item and the probe.

### Parameter exploration

The STSP mechanism underlying the model’s working memory performance is affected by two synaptic time constants: *τ*_*D*_ and *τ*_*F*_. Synaptic efficiency in the model is determined by the combination of available resources (reflecting neurotransmitters) and presynaptic calcium levels. *τ*_*D*_ determines the speed of the replenishment of the resources, while *τ*_*F*_ sets the decay rate of the calcium levels. We set these parameters to the values used in the Mongillo model (200 and 1500 ms, respectively; [12]), who, in turn, based these values on measurements in ferrets by Wang and colleagues [21]. To test how robust our model is with respect to these parameter choices, we systematically varied *τ*_*D*_ and *τ*_*F*_ over an interval of 100-400 ms and 600-1800 ms, respectively, encompassing both the range reported by Wang and the values used by Mongillo.

Figure 7A shows the effects on resource and calcium levels, as well as on the representations, split into four parameter quadrants (cf. Fig 3). The figure illustrates that higher *τ*_*F*_ values result in longer lasting elevated calcium levels and better memorization. In addition, lower *τ*_*D*_ values result in higher average spike rates (as resources are quicker replenished), also increasing calcium levels. Consequently, a combination of high *τ*_*F*_ values with low *τ*_*D*_ values results in optimal memorization and performance. This is further illustrated in Figure 7B, which shows the behavior of the model for the different parameter combinations, with the human data superimposed and associated R^2^-values and root-mean-square deviations. While correlations stay high across almost the complete parameter space, the RMSD values show that *τ*_*F*_ should be at least one second, and *τ*_*D*_ 300 ms or less for human-like behavior.

**Fig 7.**
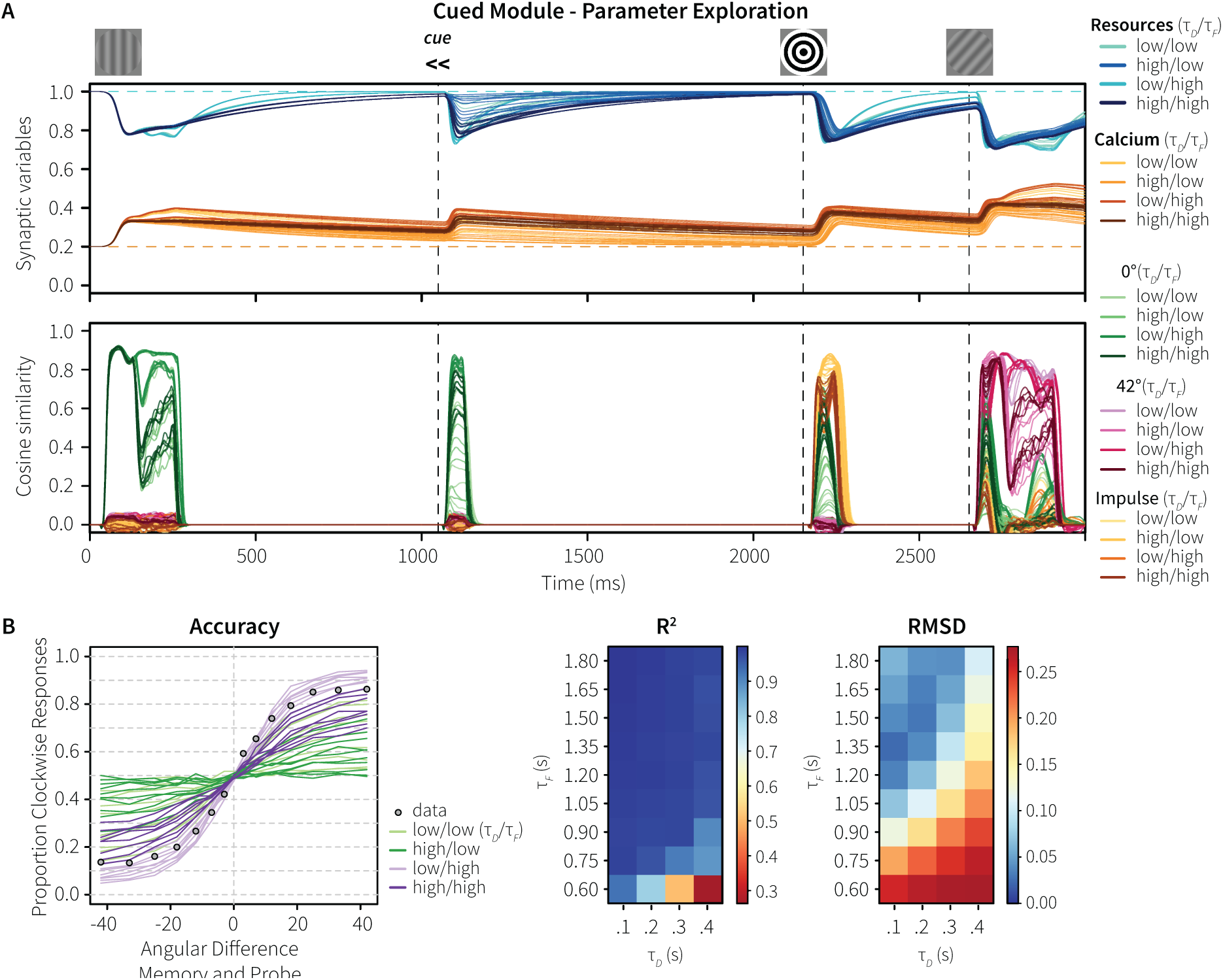
Parameter exploration of Experiment 1. A, top: resource and calcium levels for four parameter quadrants. A, bottom: absolute normalized cosine similarity between the vector represented by the memory populations and ideal vectors for 0° and 42° gratings and the impulse. All values are averaged over 100 trials with 0° memory items and 42° probes with constant within-trial phase. B: Accuracy for four parameter quadrants in comparison to the human data, and associated R2 and root-mean-square deviation values.

### Generalization 1: Longer impulse durations

To evaluate whether the model generalizes to other experimental setups, we used the same model to simulate Experiments 2 and 3 of [3], using the standard parameter values from [12]. We first discuss Experiment 3, in which Wolff and colleagues tested whether the impulse stimulus altered mnemonic representations in addition to revealing hidden states. In particular, they were concerned that the impulse stimulus would benefit behavior by reactivating the memory item. In contrast, we were concerned that the impulse stimulus disrupted the represented memory item by changing the connectivity of the recurrent memory connections through the STSP mechanism, decreasing task performance.

To test this, the design of Experiment 1 was adapted slightly (Fig 8A). The impulse stimulus could be presented at a stimulus-onset asynchrony (SOA) of 500, 250, 100, 50, or 0 ms before the probe. At the same time, the impulse always lasted until the appearance of the probe – thus the duration of the impulse varied between 500 and 0 ms. The total period between the cue and the probe was kept the same in all conditions (1400 ms, as in Experiment 1). Thus, in longer SOA conditions, the duration of the impulse was also longer, potentially affecting the memory representation more. Finally, to reduce eye strain and forward-masking effects, the bull’s eye impulse was replaced by a completely white impulse stimulus [3].

**Fig 8.**
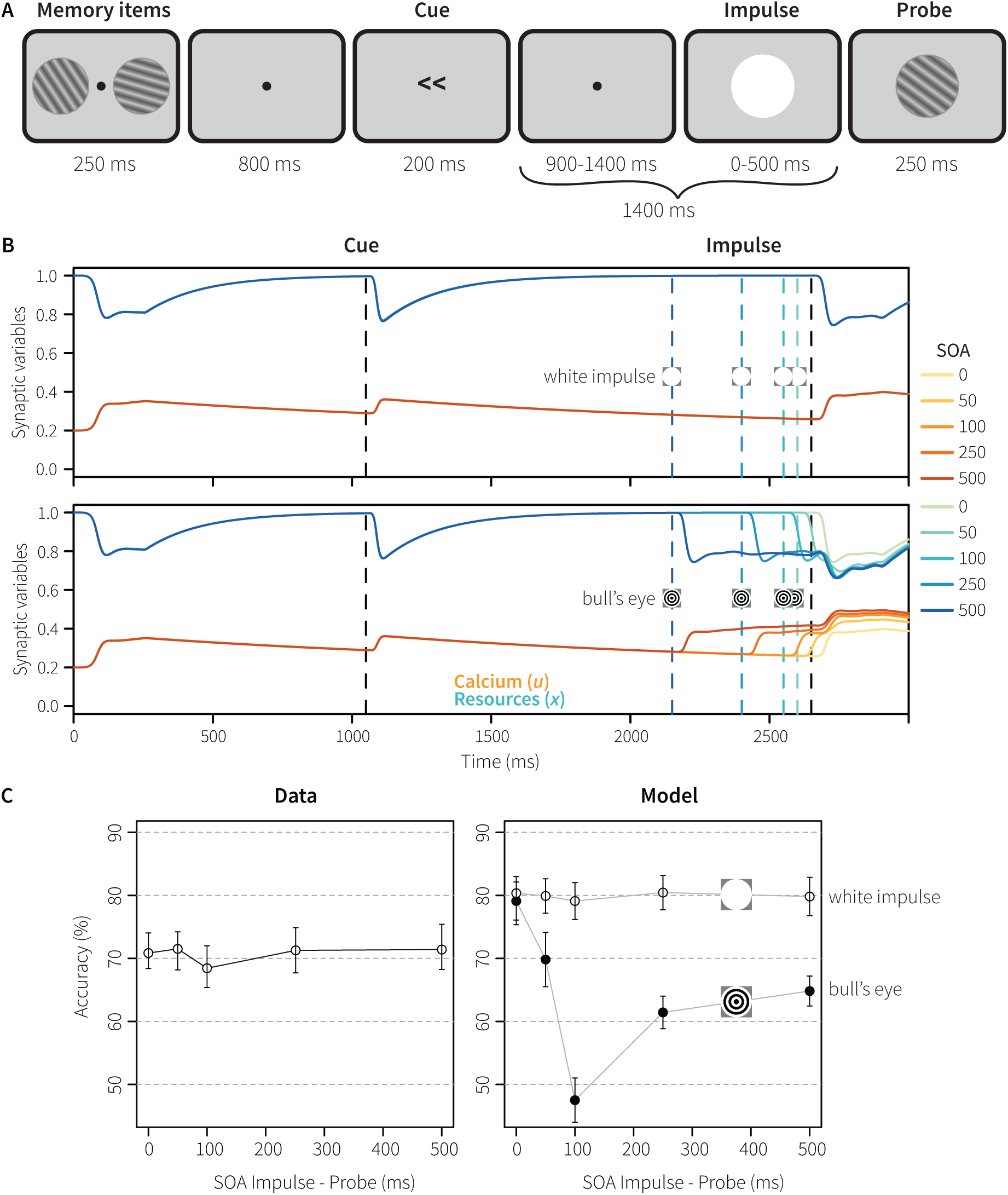
Generalization 1: Experiment 3. A) experimental procedure in which the impulse was presented at five different SOA’s and corresponding durations. B) associated calcium and resource levels. C) accuracy at the probe; angular difference between memory item and probe was always ±16 degrees.

Figure 8B shows the calcium and resource levels, Figure 8C the behavioral results. The duration of the impulse neither increased nor decreased performance for the participants (Fig 8C, left). In our first simulation with a white impulse, the model showed a similar null-effect, except that it performed about 10% better overall (Fig 8C right). On closer inspection, it transpired that a white impulse stimulus hardly reactivated the model’s memory neurons, as it hardly caused a reaction in the sensory population due to its use of Gabor filters as encoders (Fig 8B). To further test the effect, we repeated the simulation with a black bull’s eye impulse as in Experiment 1. As illustrated in Fig 8, the black bull’s eye caused a dip in performance, in particular at an SOA of 100 ms. However, with longer SOA’s performance recovered. The dip at an SOA of 100 ms is caused by a depletion of the resources of the memory neurons, decreasing their reactivation in reaction to the probe (including a slight delay)which is sufficient to make comparison and decision more difficult. With longer SOAs, the resources have been partly replenished at the moment of the probe. With a shorter SOA of 50 ms, the memory neurons are not depleted yet, and are partly activated by both the impulse and the probe. Interestingly, the original human also data suggest a similar (non-significant) dip at an SOA of 100 ms.

### Generalization 2: Maintaining multiple memory items

In Experiment 2, Wolff and colleagues investigated the maintenance of two memory items. One of the items was designated as the primary item that would be tested first, while the other item would be tested later. Participants were aware of this difference. Between the memory items and both probes white impulse stimuli were presented (Fig 9). To simulate this and generate sufficient activation after the impulse stimuli we used a 60%-grey bull’s eye for the model. We used a 60%-grey value as it led to clearly visible effects for illustrative purposes, as well as resulting in reasonable behavior. However, this is an arbitrary choice and does not affect the qualitative effects (see also the Discussion).

**Fig 9.**
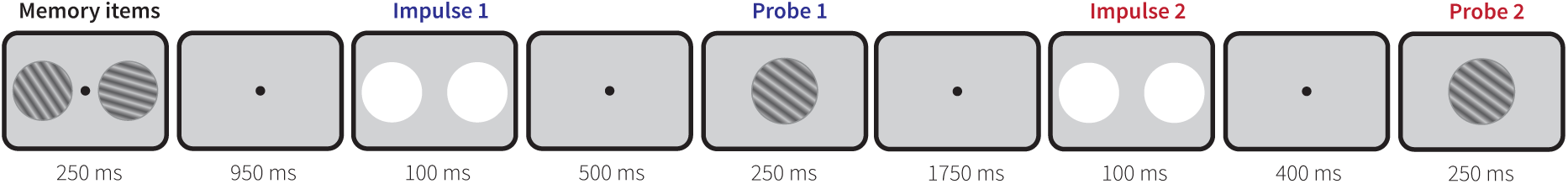
Procedure Generalization 2/Experiment 2. Participants knew which memory item would be probed first. Note the long break after Probe 1.

In reaction to the first impulse, both memory items could be decoded, but the primary item was slightly better than the secondary item – indicating an attentional difference [3]. To simulate this we gave the sensory module of the primary item the same input as in the previous simulations, while the sensory module reflecting the secondary item received 90% of the input. About 450 ms after the onset of the first probe, the onset of a strong lateralization in the EEG signal was observed, similar to the effect of the cue in Experiment 1. We therefore simulated this likewise with a non-specific population-wide input to the memory module of the secondary memory item, reactivating its representation, and assuming participants similarly refreshed the secondary memory item after making the first response. Otherwise the model was identical to the simulations described above.

Fig 10 shows the spiking activity and representations over the course of a trial (A: primary memory item, probed early; B: secondary item, probed late). As in the simulation of Experiment 1, the representations could not be decoded in between the various stimuli, indicating activity-silent storage. The reaction to the initial presentation of the memory items is a little stronger for the primary than for the secondary item, reflecting the input difference to the sensory modules. This is further reflected in response to the first impulse, which is higher for the primary item. At 1800 ms, the first probe is used to make a decision for the primary item, assuming that participants could flexibly gate this to the correct module (not included in the model, but see [34] for a potential implementation). The secondary memory module is reactivated 450 ms later with a population-wide input: the resulting activation clearly reflects the original stimulus. After a long pause (1750 ms), both modules again receive an impulse stimulus, followed by the probe for the secondary memory item.

**Fig 10.**
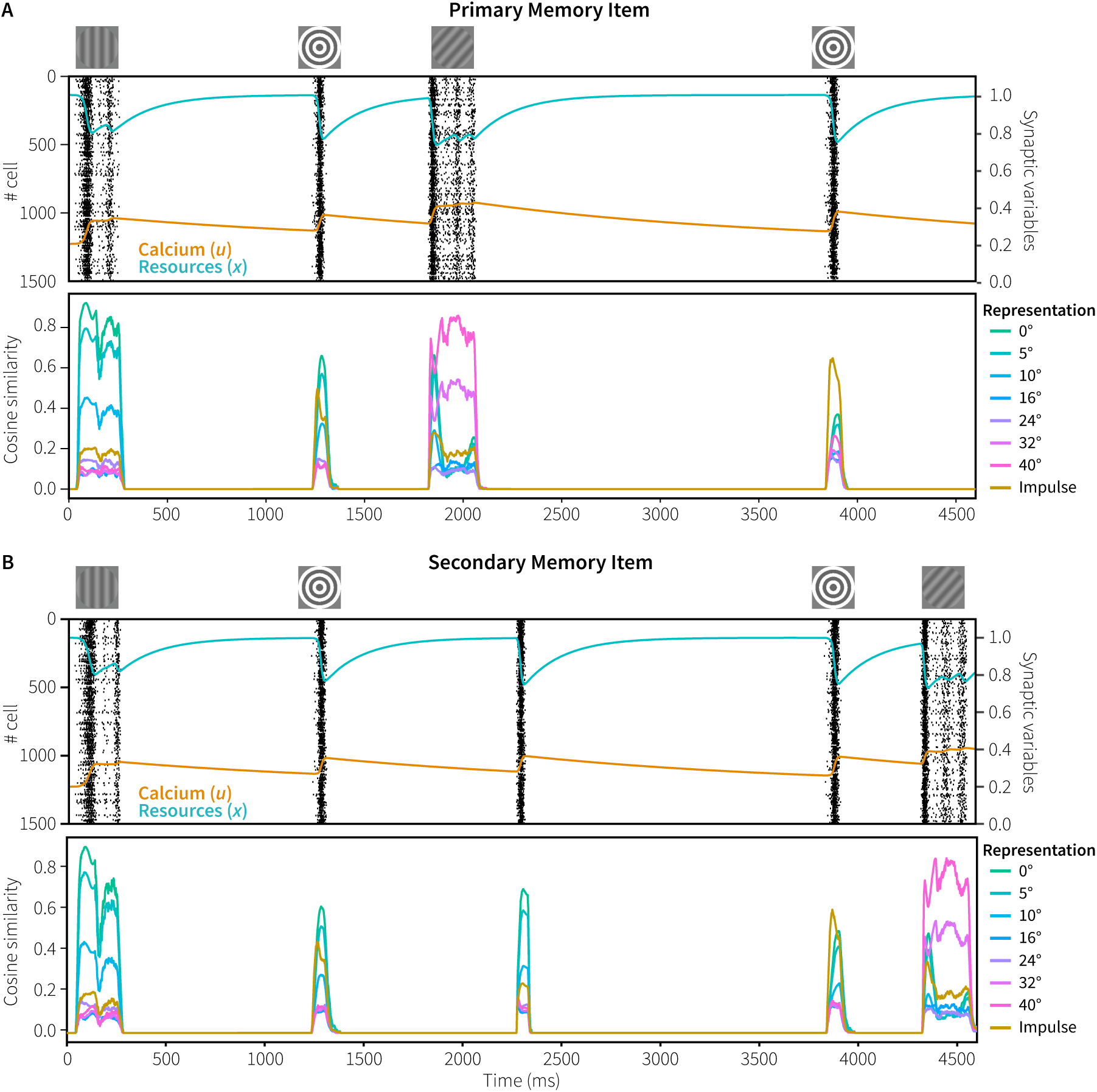
Spiking behavior and representations in Generalization 2/Experiment 2. A) primary memory item. B) secondary memory item. Representations averaged over 100 trials with 0° memory items and 40° probes.

Fig 11 shows the reaction to both impulse stimuli in more detail. First, and somewhat trivially, the reaction to the impulses is weaker than in Experiment 1, because we used a 60% grey bull’s eye instead of a black one. Second, in response to the first impulse both items could be decoded in the experiment, but the primary item a little better [3]. This is reflected in the model’s results, where the difference between the impulse representation and the memory item is larger for the primary item. Third, in reaction to the second impulse, only the secondary item could be decoded in the experiment. The model also reflects this, as the representation of the primary item is about half of the representation of the impulse, while the secondary item just exceeds the impulse (cf. Fig 4).

**Fig 11.**
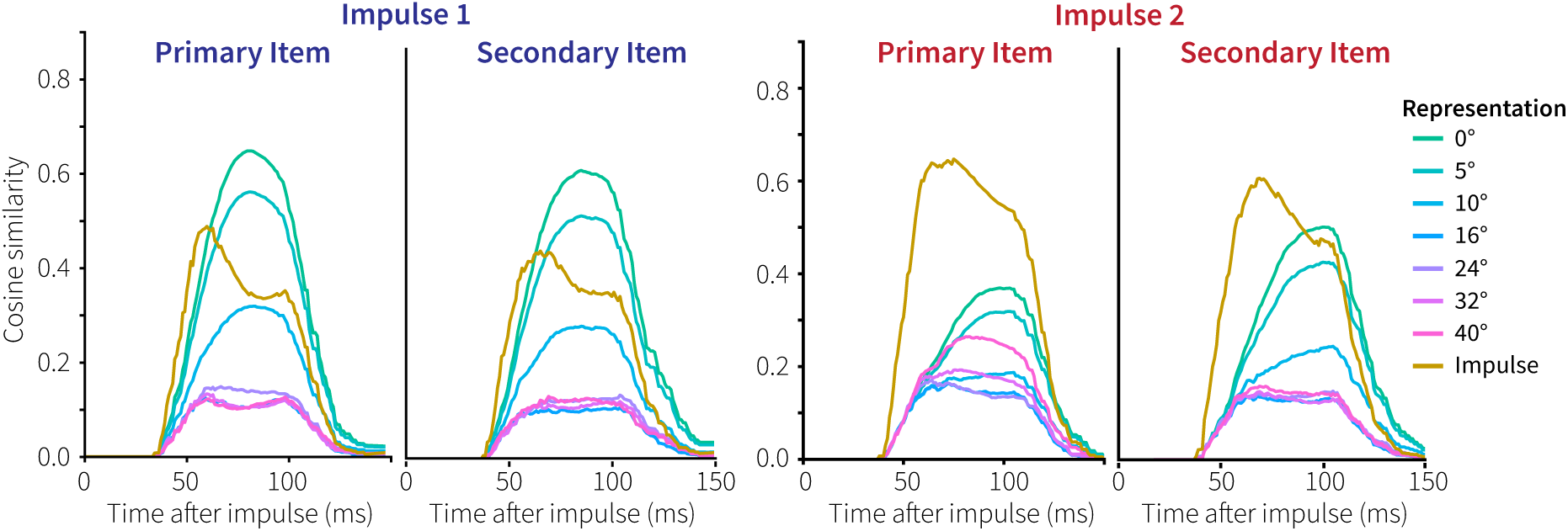
Primary and secondary memory representations in response to the impulse stimuli in Generalization 2/Experiment 2.

Finally, as one of our main goals was to simulate human behavior, Fig 12 compares the model’s performance to the human participants. The model performs slightly better on the largest angular differences for the primary memory items than participants, but overall the correspondence to the data is remarkable. The decreased performance to the secondary memory item is due to the slightly lower input to the sensory module at the start of the trial, and to the much longer interval between the reactivation and the probe. However, reactivating the item once was sufficient for human-like performance, suggesting that no rehearsal took place during the maintenance interval.

**Fig 12.**
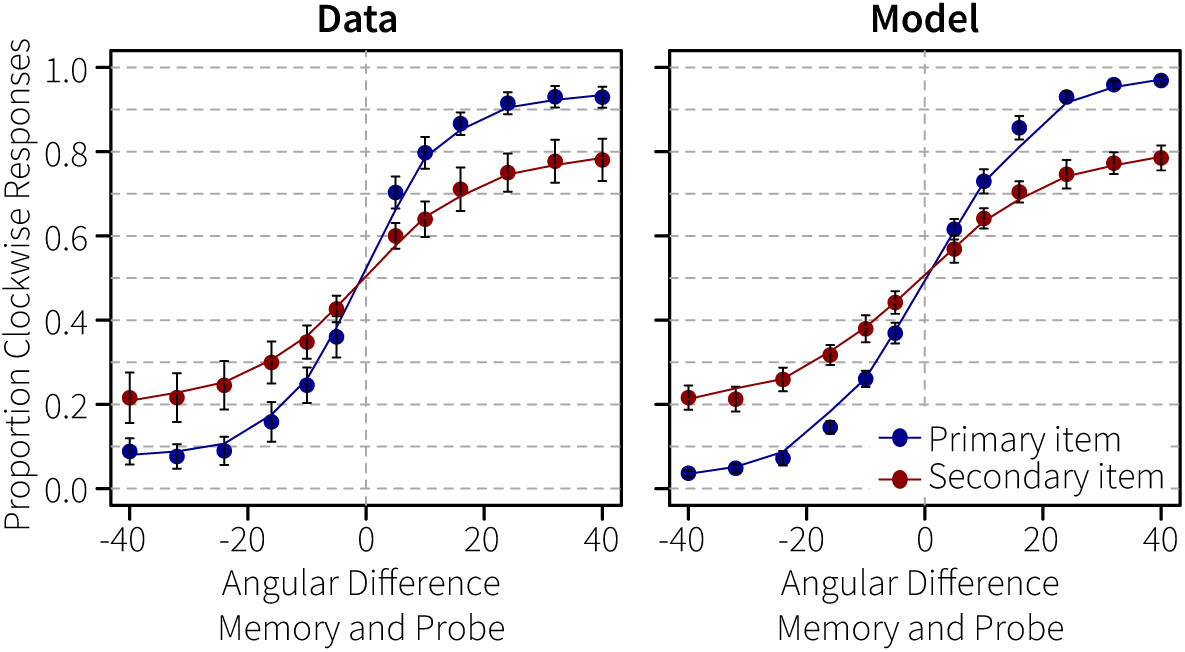
Performance in Generalization 2/Experiment 2. Proportion of clockwise responses dependent on the angular difference between the memory items and the probes.

## Discussion

We developed a functional spiking-neuron model to explain recent theories of activity-silent human working memory. Whereas incoming information is encoded through spiking, maintenance of information was realized by short-term synaptic plasticity based on calcium kinetics [12]. This mechanism can maintain information effectively for short periods of time without requiring neural spiking. In support of the model, we simulated a recent EEG experiment from a study that applied an innovative impulse perturbation method [3] to reveal the content of activity-silent WM. Both the model’s choice behavior, as well as its mental representations corresponded well to the human data.

To further characterize the model and test its generalizability, we subsequently performed a parameter exploration and simulated two additional experiments. First, the parameter exploration showed that the model’s behavior is robust over a range of parameter values, unless the decay rate of calcium, *τ*_*F*_, goes below one second. In that case performance breaks down, implying that the calcium decay rate in human working memory regions either exceeds one second (which is possible given the reported values in ferrets, see [21]), or that a reactivation mechanism is used for maintenance over longer periods (see below). Second, it was shown that the activity-silent memory representations are not very sensitive to the impulse response, matching human data. Thirdly, the model could mimic memorizing two stimuli that were probed sequentially. Also in this case the mental representations of the model seemed to be in agreement with the human data, maintaining *and* forgetting information at the same rate as human participants. Taken together, this demonstrates that calcium-mediated STSP not just results in robust maintenance of arbitrary stimuli, as shown earlier [12], but can also simulate effective human behavior based on real-world stimuli.

With regard to localization, the model was used to simulate data from Wolff and colleagues [3], who reported posterior EEG effects. However, WM is often attributed to prefrontal areas [e.g., 14,35]. Activity-silent maintenance has likewise been found in both posterior [3,8,36] and frontal [18,37] regions. It appears that especially sensory working memory should be attributed to the relevant sensory systems [36,38], instead of to a centralized system. While the exact function of the different regions implicated in WM might differ, the neural substrate and mechanisms might be similar, and could potentially all be explained by the proposed STSP mechanism.

A number of design choices warrant discussion. First, the employed neurons do not have a baseline firing rate, as is evident by the lack of any spiking activity during the delay-period of a trial (Fig 3). In order to clearly demonstrate activity-silent maintenance of information, we defined the tuning curves of the sensory and memory neurons so that they only fire when presented with input. As a result, the memory was fully silent when no input was presented. If one would broaden the tuning curves of the neurons, the recurrent connection would remain active after having received input, resulting in activity-based working memory. On the other hand, background firing could simply be added to the model without affecting its functionality, as has been done in the past ([e.g., 12,34]; see also below for an example). Second, the number of neurons per population and the number of dimensions used to represent the stimuli were set to reflect human behavior. In general, adding more neurons will improve the representation of vectors and the approximation of the functions computed over those. Increasing the number of dimensions expands what can represented [26,39,40]. Thus, changing the number of neurons and dimensions will change the quality of the representations and will influence the number of errors made during the task. Here, we estimated parameters to roughly match human performance; we do not have a principled reason either for using 1000 or 1500 neurons per population or 24 dimensions.

For our second generalization, Wolff’s Experiment 2, we used a 60%-grey bull’s eye instead of the completely white impulse stimulus that was used in the actual experiment. We chose 60%-grey to ensure sufficient activation in response to the impulse stimuli both for illustrative as well as behavioral purposes, and because we had seen in our first generalization that the model’s memory neurons were hardly reactivated by a white impulse. We believe that this is due to the very simple visual system (i.e. a single layer of Gabors), and does not affect the core results. As the visual system is not the focus of the current modeling effort, we decided to solve this by using a grey-scale version of the bull’s eye instead of using a more complex visual system. As mentioned above, 60% grey was an arbitrary choice, as any bull’s eye stimulus with sufficient contrast yields the same qualitative effects.

Finally, information was represented using Nengo’s default vector representation, which provides an intuitive method to link neural spiking to representation and function [26]. However, representing information differently should not affect the basic functioning of the model as all connections and the STSP mechanism are implemented at the neural level. Thus, while Nengo made implementing a real-world task straightforward, the mechanisms that we used are general and could be used in any framework.

### Representations in WM

As discussed above, in the current model information is maintained without *any* intermittent firing (Fig 3). This directly contradicts the original analysis of the dataset [3], where the represented stimulus could be decoded for some time after its offset. In addition to fully activity-silent maintenance, Mongillo and colleagues [12] observed a bi-stable regime in their model: with added background noise, neurons with facilitated connections reactivated spontaneously. Consequently, due to the dynamics of *u* and *x*, the reactivated neurons will be briefly depressed before being facilitated again, leading again to reactivation. In this regime, the time between subsequent reactivations is on the scale of *τ*_*D*_ – the time constant of the available resources – as it is controlled by the recovery from the synaptic depression. A brief exploratory analysis shows that such a bi-stable regime can also be added to our model, as is illustrated in Fig 13. This provides the model with an additional method of maintaining information, possibly over a longer period of time. In case human calcium decay rates are in fact below one second, as discussed above, this might also be the way in which the human brain stores information during the delay periods of the current experiments. Finally, it provides a potential explanation for the delay-activity observed in the original analysis [3]: non-specific background or recurrent input after presentation of a stimulus might temporarily have pushed the network into this regime.

**Fig 13.**
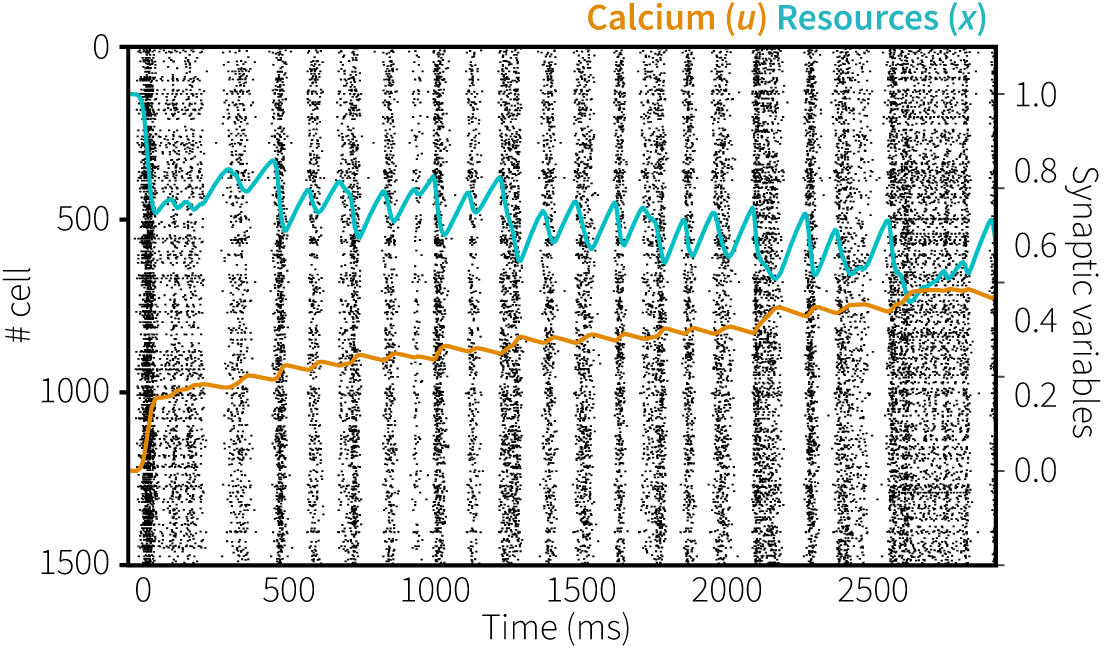
Bi-stable maintenance. Background noise puts the model in a bi-stable regime where facilitated connections reactivate spontaneously on the time scale of *τ*_*D*_.

Another functional role for delay activity in WM might be tracking the focus of attention [15–17,41]. This would provide a mechanistic explanation for psychological theories that state that a single focal WM item can be used without any time cost [16,42–44], while other items in working memory incur a cost estimated at 200 ms [15,45–48] – the latter potentially being due to the costs of reactivating the information from a non-active state. Following previous conceptions [41,49], Wolff and colleagues [3] suggested that a difference in focal attention might also dissociate the maintenance of the primary item and the secondary item in Experiment 2. Here, we have shown that we can explain their behavioral data more simply by assuming a slight attentional difference at encoding and an attentional pulse for the secondary item following the first probe. However, they showed slightly elevated delay activity for the primary memory item, suggesting that the primary item in the focus of attention is not stored in a completely activity-silent state. In the interest of simplicity, this is not reflected by the current model.

Wolff and colleagues further suggested that their experiments show that information could be quickly and flexibly cleared from memory. As decoding the uncued stimulus in Experiment 1 and the primary memory item at the second probe in Experiment 2 was not possible, it was argued that these must have been deleted. However, our models show that this is does not necessarily follow. As a result of not reactivating the uncued item at the cue in Experiment 1, it had only half the representational power of the impulse at the moment of the impulse (Fig 4, right). Similarly, in Experiment 2 the combination of the first probe overwriting the primary item and the long delay to the second impulse results in the original memory item only being very weakly represented (Fig 11, third panel). Thus, although the memory items might have been cleared actively, these experiments do not seem to require that, nor do we see a straightforward way of adding such a mechanism to the model.

As implied above, we assume that the memory vector being only half the strength of the represented impulse means that the memory is effectively forgotten (unless the same stimulus is presented again). At least, we assume it will be hard to decode from the other representations in the EEG signal, and therefore argue that our models represent the decoding results presented in [3]. However, one can wonder how sensitive this effect is to our choice of parameters. The effect is based only on the parameters of the STSP mechanism, which we took directly from [12], in combination with the timing of the experiment. In this sense it is a parameter-free model fit, which turns out to work well. Our parameter exploration, in particular Fig 7A, shows that different values for *τ*_*F*_ and *τ*_*D*_ lead to different representation ratios between the represented memory items and the representation of the impulse. However, the qualitative effects – the difference between uncued and cued items – remains constant, until *τ*_*F*_ becomes too low and behavioral performance decreases. It does predict that if we reduce the time between the memory items and the impulse, decoding the item from the uncued model should be possible. While not tested directly, Experiment 2 did present an impulse after about 1 second (instead of a cue), after which both items were decodable. In addition, after the cue in Experiment 1, both items could still be decoded [Supplementary Fig 1 of 3].

In spite of these effects of time on the uncued item, the represented memories were remarkably resistant against corruption by longer impulses, as indicated by the simulation of Experiment 3. One reason for this is a quick depletion of the resources of the neurons, leading to a strong reduction in overall spiking and therefore to less disruption of the original representation. This could simply be a result of the calcium-based STSP mechanism, or it might be a functionally relevant protection mechanism, resulting in more stable representations.

In our model, we have assumed that the coding of the information itself is static, that is, the same facilitated connections are used repeatedly. However, there has recently been increasing evidence for a dynamic coding framework, which states that information maintained in a WM network traverses a highly dynamic path through neural activation [37]. It is not yet clear how this relates to our model, although a possible clue might come from a model by Singh and Eliasmith [50]. Neural populations in their model represent two dimensions, where one dimension represents time and the other a stimulus. Because their neurons used tuning curves sensitive to both dimensions, the neural responses also changed continuously – resulting in a dynamic representation. Their model elegantly captures a wide variety of observed neural responses during a WM task; the inclusion of time as a dimension represented by the neurons in the network naturally leads to a dynamic firing pattern over time. The current model could likewise be adapted to also represent time, but as we have no clear evidence that this is the case, we decided against it. We do plan to explore this in future work.

### Related Models

Recently, Myers and colleagues [8] described a related non-spiking neural population model with similar functionality as the current model, although they did not match human data directly. Their model consisted of a three-layer architecture: a stimulus layer, a template layer, and a decision layer not unlike the sensory, memory, and comparison population in our model. A critical difference between the two models is that their decision layer only receives input from the template layer, while in our case it receives input from both the sensory and memory populations. The template layer in Myers’ model acts like a match-filter: it is able to maintain a stimulus orientation, and when presented with a subsequent probe orientation convey the signed difference between the two to the decision layer. The memory population in our model can likewise be viewed as a match filter. After onset of the probe, the represented orientation shifts to the orientation of the probe from the direction of the orientation of the memory item. This shift in itself indicates a degree of difference between the two orientations, including the sign of this difference. One could potentially measure this with a neural population that computes a time derivative with respect to the orientation [51]. However, exploratory analysis indicated this to be less robust than our current method.

Another closely related model was proposed by Barak and colleagues [18]. Their model consisted of a sensory and memory population. After presentation of a stimulus, connections from the sensory population to the memory population will be facilitated. Subsequently, during the delay period, an increasing uniform current is applied to the network which activates the neurons in the memory population that have facilitated incoming connections. During presentation of a subsequent probe, mutual inhibition between the sensory and memory population will guide decision making. This model explains observed ramping up of activity during anticipation of a probe. However, it is not clear whether the gradually increasing external current is essential to extract the information maintained in the facilitated connections in the memory representation. It can be expected that in the brain bottom-up stimulus driven activity might also be able to activate the information stored in connections, for instance when the timing of the probe is unknown.

### Conclusion

To conclude, our model shows that maintenance of information in WM by means of calcium-mediated STSP can lead to functional behavior. It is broadly consistent with current theories regarding activity-silent storage in human WM and is able to show a variety of effects observed during three visual delayed-response tasks. Furthermore, it provides a solid basis for exploring a model that incorporates psychological theories on the focus of attention [15–17,42] by combining activity-silent maintenance with storage through persistent firing.

## Methods

### Model

#### Nengo

The model was implemented using Nengo, a Python library for simulating large-scale neural models with a clear link between spiking activity and representation [25–27]. Nengo makes use of a theoretical framework called the Neural Engineering Framework [NEF; 52]. Information is represented as a vector of real numbers that can be encoded and decoded from the collective spiking activity of populations of neurons. Encoding is mediated by giving each neuron a non-linear tuning curve that characterizes their general response to the incoming signal. Decoding is a linear process: the activity of each neuron in a population is weighted by a constant and summed over time in order to decode the represented vector. Connections between populations allow for the communication and transformation of the information. Here the NEF calculates the connection weight matrix between populations to approximate a desired function. In addition, connection weights can be learned and adapted through several biologically plausible learning rules, both supervised and unsupervised [28].

#### Short-term synaptic plasticity

Short-term synaptic plasticity was implemented in Nengo following the calcium kinetics mechanism of Mongillo and colleagues ([12]; available at https://github.com/Matthijspals/STSP). Because spiking leaky integrate-and-fire (LIF) neurons are computationally efficient while retaining a degree of biological plausibility, we added this mechanism to the existing Nengo implementation of LIF neurons. Synaptic efficiency is based on two parameters: the amount of available resources to the presynaptic neuron (*x*, normalised between 0 and 1) and the fraction of resources used each time a neuron fires (*u*), reflecting the residual presynaptic calcium level. When a neuron fires, its resources *x* are decreased by *ux*, mimicking neurotransmitter depletion. At the same time, its calcium level *u* is increased, mimicking calcium influx into the presynaptic terminal. Both *u* and *x* relax back to baseline with time constants *τ*_*D*_ (0.2s) and *τ*_*F*_ (1.5s), respectively. This results in a system where after a neuron fires its outgoing connections will be depressed on the time scale of *τ*_*D*_ and facilitated on the timescale of *τ*_*F*_ as illustrated in Fig 14 (darker lines indicate standard parameter values, lighter lines the range tested in the parameter exploration).

**Fig 14.**
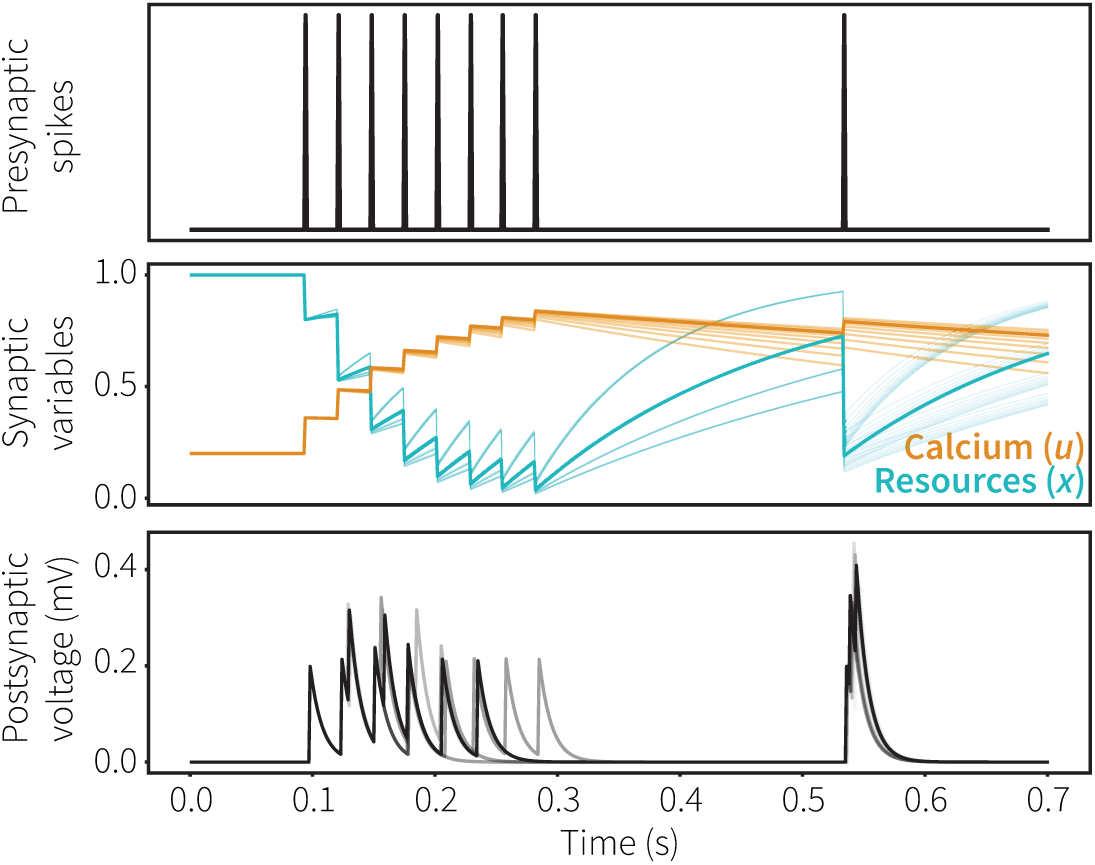
STSP mechanism. Top: spikes in presynaptic neuron. Middle: calcium (*u*) and resources (*x*) of presynaptic neuron, *u* increases and *x* decreases when the presynaptic neuron spikes. Bottom: resulting postsynaptic voltage; note the synaptic depression at the end of the first spike train and synaptic facilitation at the later spike. Dark lines indicate standard parameter values, lighter lines different parameter settings for *τ*_*D*_ (100-400 ms) and *τ*_*F*_ (600-1800 ms). Nengo reproduction of Fig 1A in [12].

For all LIF neurons to which we apply STSP, every time step *u* and *x* are calculated according to Equation 1.1 and 1.2, respectively:

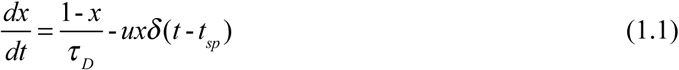

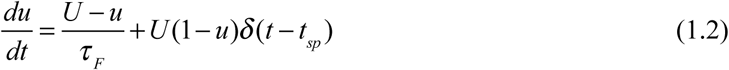

where *x* represents the available resources, *u* the residual calcium level, *τ*_*D*_ the depressing time constant, *δ* the Dirac delta function, *t* the simulation time and *t*_*sp*_ the time of a presynaptic spike. In equation 1.2, *τ*_*F*_ represents the facilitating time constant and *U* the calcium baseline level. Outgoing connection weights of neurons implementing STSP are determined by both their initial connection weight and their current synaptic efficiency. Initial connections weights are calculated by the NEF, while synaptic efficiency is set to the product of the current value of *u* and *x* of the presynaptic neuron, normalised by their baseline value:

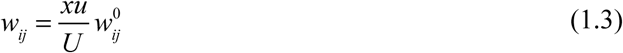

where *w*_*ij*_ represents the connection weight between neuron *w*_*ij*_^0^ the initial connection weight.

#### Architecture

The overall architecture of the model is shown in Fig 2 (the model is available for download at https://github.com/Matthijspals/STSP). The sensory and decision populations consist of 1000 LIF neurons, the memory and comparison populations of 1500 LIF neurons. Biologically relevant parameters were left to default, which are consistent with neocortical pyramidal cells [25]. Parameters *U, τ*_*D*_ and *τ*_*F*_ were set the same as in [12], except in our parameter exploration. *τ*_*F*_ ≫*τ*_*D*_ and *τ*_*F*_ on the order of 1s are consistent with patch-clamp recordings of facilitated excitatory connections in the ferret prefrontal cortex [21]. Table 1 lists all parameter settings.

**Table 1.** Model parameters for each population. Default parameters in italics.

| Parameter | Sensory | Memory | Comparison | Decision |
| --- | --- | --- | --- | --- |
| Number of neurons | 1000 | 1500 | 1500 | 1000 |
| Neuron type | LIF | STSP-LIF | LIF | LIF |
| Dimensions | 24 | 24 | 4 | 1 |
| Intercepts | Uniform(0.01, .1) | Uniform(0.01, .1) | Uniform(.01, 1) | *Uniform(-1, 1)* |
| $\tau_{rc}$ (Membrane RC) | *0.02* | *0.02* | *0.02* | *0.02* |
| $\tau_{ref}$ (refractory period) | *0.002* | *0.002* | *0.002* | *0.002* |
| $U^*$ | | 0.2 | | |
| $\tau_D^*$ | | 0.2 | | |
| $\tau_F^*$ | | 1.5 | | |

\*taken from Mongillo, Barak and Tsodyks (2008)

To describe the relationship between neural representations and real-world stimuli it can be assumed that the brain makes use of a statistical model, not unlikely a parametrized model, where a small number of parameters capture the overall shape of the data [26]. Ensembles of neurons in a Nengo model represents information as a *D*-dimensional vector of real numbers. We can represent the stimuli in our experiment by finding *D* parameters that best represent these stimuli and letting these be the values that our neural ensembles represent.

To find these parameters we need a set of *D* basis functions that will be good at describing both the incoming images and the encoders of the neurons receiving these images. These basis functions can be found by applying singular value decomposition (SVD) to a matrix *T*, containing both the images and the encoders. The images consisted of the stimuli in the experiment, while the encoders were two-dimensional Gabor filters, defined by a sinusoidal plane wave multiplied by a Gaussian function. Gabor filters have previously been shown to accurately describe the response profile of simple cells in the cat [30] and macaque [29] striate cortex and seem to underlie early stages of visual processing. Thus, the SVD mediates a biologically plausible method that results in stimuli being represented by *D*-dimensional vectors. In our model, we set *D* to 24.

Applying SVD on *T* gives us matrices *U, S*, and *V*, for which:

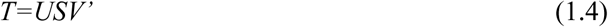

where *U* and *V* are respectively the left and right singular matrices and *S* is the diagonal matrix containing the singular values of *T*. Since the values in *U, S* and *V* are ordered by how much they contribute to *T*, we can represent *T* reasonably well by just taking the top singular values (similar to principle component analysis, PCA); in order to transform input into a *D* dimensional vector that can be represented by our neurons, we used the top 24 singular values. This allows us to specify our encoders *E* as follows:

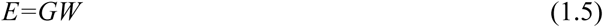

where *G* are the generated Gabor filters and *W* the top *D* columns of the matrix *U*. Any input in the network that has passed through these encoders will be represented by a vector of length *D*.

Now that input is represented by a vector of length *D*, we need an ensemble that takes this vector and decodes it into orientations. As a consequence of how we specified the encoders, the *D* dimensional matrix *R* the stimuli will be represented by is equal to:

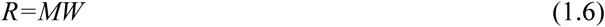

where *M* are the possible stimuli and *W* the top *D* columns of the matrix *U*. Next, we specified for each stimulus-vector – representing a particular grating – what the corresponding decoded orientation should be. This information was used to define a function that takes a vector as input and returns the corresponding orientation. The NEF then yields the connection matrix at the neural level that approximates this function. This matrix was used to set the connections from the sensory and memory populations to the comparison populations.

Stimulus orientations were not directly decoded as the angle *θ*, but rather by the sine and cosine of *θ*. Decoding sine and cosine of *θ* is robust, as the ratio between the two determines the stimulus orientation independent of the amplitude, which is not the case when decoding *θ* directly. Furthermore, the symmetry of the sine functions provides a natural solution for the symmetric nature of the stimuli, as in the experiment a stimulus with an orientation of −90° contains exactly the same pixels as a stimulus with an orientation of 90° and therefore results in the same neural activity.

### Experimental Simulation

#### Stimuli

Input to the model consisted of images of 128 by 128 pixels. Stimuli were generated using Psychopy, an open-source Python application [53]. Stimuli consisted of a circle on a grey background (RGB = 128, 128, 128). Memory items and probe stimuli were sine-wave gratings with a diameter of 128 pixels and spatial frequency of 0.034 cycles per pixel. The phase was randomized within and across trial. For each trial, the orientation of the memory items was randomly selected from a uniform distribution of orientations. In Experiment 1, the angular differences between the memory item and the corresponding probe stimulus were uniformly distributed across seven angle differences (3°, 7°, 12°, 18°, 25°, 33°, 42°), both clockwise and counter-clockwise. The impulse stimulus consisted of a black ‘bull’s-eye’ stimulus of the same size and spatial frequency as the memory items. It was presented at twice the contrast compared to the grating stimuli, to each module.

In Generalization 1 (Experiment 3 in [3]), the angular difference between the memory item and the corresponding probe stimulus was always 16°, clockwise and counter-clockwise. The impulse stimulus was either a white circle or the black ‘bull’s-eye’ stimulus from Experiment 1.

In Generalization 2 (Experiment 2 in [3]), the angular differences between the memory item and the corresponding probe stimulus were uniformly distributed across six angle differences (5°, 10°, 16°, 24°, 32°, 40°), both clockwise and counter-clockwise. The impulse stimulus consisted of a 60% grey ‘bull’s-eye’ stimulus of the same size and spatial frequency as the memory items. It was presented at twice the contrast compared to the grating stimuli, to each module.

#### Procedure

The model simulated the three the retro-cue delayed-response tasks from [3]. In Experiment 1, each trial started with the presentation of two memory items to the sensory population of the corresponding modules for 250 ms. In the original experiment, a retro-cue that indicated which of the two previously presented items needed to be memorized was presented 800 ms later, which was followed by significant lateralization at posterior electrodes. To mimic this, the memory population of the cued module is briefly reactivated by means of a non-specific population wide input with an amplitude of 0.02 for 20 ms [cf. 12]. After another fixation period, the impulse stimulus was presented to both sensory populations for 100 ms, 1100 ms after the onset of the cue. After another delay of 400 ms, the probe was presented to the sensory populations for 250 ms. To simulate different participants in the experiment, every 1344 trials the random seed was reset and new random Gabor filters were generated to use as encoders for the sensory populations. In total the model performed 30 sets of 1344 trials, reflecting 30 participants in the original experiment.

Generalization 1 (Experiment 3 in [3]) was identical to Experiment 1, except that the impulse stimulus was presented at 5 different SOA with respect to the probe (0, 50, 100, 250, 500 ms). In addition, the impulse stimulus now lasted until the probe was presented. In the 0-ms case, no impulse stimulus was shown. To simulate different participants in the experiment, every 280 trials the random seed was reset and new random Gabor filters were generated to use as encoders for the sensory populations. In total the model performed 20 sets of 280 trials, reflecting 20 participants in the original experiment.

In Generalization 2 (Experiment 2 in [3]), each trial started with the presentation of two memory items to the sensory population of the corresponding modules for 250 ms. After a fixation period of 950 ms, the first impulse was presented to both sensory populations for 100 ms. After another fixation period, the first probe was presented to the sensory population of the primary memory item, for 250 ms. To mimic reactivating the secondary memory item afterwards, the cued module is briefly reactivated by means of a non-specific population wide input with an amplitude of 0.02 for 20 ms, 450 ms after the onset of the first probe (as indicated by the lateralization in the EEG data, see the main text and [3]). 1750 ms after the first probe the second impulse was presented to both sensory populations for 100 ms. After another delay of 400 ms, the second probe was presented to the sensory population of the secondary memory item for 250 ms. To simulate different participants in the experiment, every 1728 trials the random seed was reset and new random Gabor filters were generated to use as encoders for the sensory populations. In total the model performed 19 sets of 1728 trials, reflecting 19 participants in the original experiment.

## References

1. Baddeley AD, Hitch GJ. Working Memory. In: Bower GA, editor. Psychol. Learn. Motiv. New York: Academic Press; 1974. page 47–89.

2. Barak O, Tsodyks M. Working models of working memory. Curr. Opin. Neurobiol. 2014;25:20–4.

3. Wolff MJ, Jochim J, Akyürek EG, Stokes MG. Dynamic hidden states underlying working-memory-guided behavior. Nat. Neurosci. 2017;20(6):864–71.

4. Rombouts JO, Bohte SM, Roelfsema PR. How attention can create synaptic tags for the learning of working memories in sequential tasks. PLoS Comput. Biol. 2015;11(3):e1004060.

5. Fuster JMJ, Alexander GE. Neuron Activity Related to Short-Term Memory. Science. 1971;173(3997):652–4.

6. Goldman-Rakic PS. Cellular basis of working memory. Neuron. 1995;14(3):477–85.

7. Lundqvist M, Herman P, Miller EK. Working Memory: Delay Activity, Yes! Persistent Activity? Maybe Not. J. Neurosci. 2018;38(32):7013–9.

8. Myers NE, Rohenkohl G, Wyart V, Woolrich MW, Nobre AC, Stokes MG, et al. Testing sensory evidence against mnemonic templates. Elife. 2015;4:1–25.

9. Sreenivasan KK, Curtis CE, D’Esposito M. Revisiting the role of persistent neural activity during working memory. Trends Cogn. Sci. 2014;18(2):82–9.

10. Watanabe K, Funahashi S. Neural mechanisms of dual-task interference and cognitive capacity limitation in the prefrontal cortex. Nat. Neurosci. 2014;17(4):601–11.

11. Lundqvist M, Rose J, Herman P, Brincat SL, Buschman TJ, Miller EK. Gamma and Beta Bursts Underlie Working Memory. Neuron. 2016;90:152–64.

12. Mongillo G, Barak O, Tsodyks M. Synaptic Theory of Working Memory. Science. 2008;319(5869):1543–6.

13. Stokes MG. ‘Activity-silent’ working memory in prefrontal cortex: a dynamic coding framework. Trends Cogn. Sci. 2015;19(7):394–405.

14. Borst JP, Anderson JR. Using Model-Based functional MRI to locate Working Memory Updates and Declarative Memory Retrievals in the Fronto-Parietal Network. Proc. Natl. Acad. Sci. USA. 2013;110(5):1628–33.

15. Borst JP, Taatgen NA, Van Rijn H. The Problem State: A Cognitive Bottleneck in Multitasking. J. Exp. Psychol. Learn. Mem. Cogn. 2010;36(2):363–82.

16. Oberauer K. Access to information in working memory: Exploring the focus of attention. J. Exp. Psychol. Learn. Mem. Cogn. 2002;28(3):411–21.

17. Olivers CNL, Peters J, Houtkamp R, Roelfsema PR. Different states in visual working memory: when it guides attention and when it does not. Trends Cogn. Sci. 2011;15(7):327–34.

18. Barak O, Tsodyks M, Romo R. Neuronal population coding of parametric working memory. J. Neurosci. 2010;30(28):9424–30.

19. Zucker RS, Regehr WG. Short-Term Synaptic Plasticity. Annu. Rev. Physiol. 2002;64(1):355–405.

20. Tsodyks M, Pawelzik K, Markram H. Neural Networks with Dynamic Synapses. Neural Comput. 1998;(10):821–35.

21. Wang Y, Markram H, Goodman PH, Berger TK, Ma J, Goldman-Rakic PS. Heterogeneity in the pyramidal network of the medial prefrontal cortex. Nat. Neurosci. 2006;9(4):534–42.

22. Jackman SL, Regehr WG. The Mechanisms and Functions of Synaptic Facilitation. Neuron. 2017;94(3):447–64.

23. Myers NE, Rohenkohi G, Wyart V, Woolrich MW, Nobre AC, Stokes MG. Testing sensory evidence against mnemonic templates. eLife. 2015;4:1–25.

24. Wolff MJ, Ding J, Myers NE, Stokes MG. Revealing hidden states in visual working memory using electroencephalography. Front. Syst. Neurosci. 2015;9(123):1–12.

25. Bekolay T, Bergstra J, Hunsberger E, DeWolf T, Stewart TC, Rasmussen D, et al. Nengo: a Python tool for building large-scale functional brain models. Front. Neuroinformatics. 2014;7:48.

26. Eliasmith C. How to Build a Brain: A Neural Architecture for Biological Cognition. New York, NY: Oxford University Press; 2013.

27. Eliasmith C, Stewart TC, Choo X, Bekolay T, DeWolf T, Tang C, et al. A large-scale model of the functioning brain. Science. 2012;338(6111):1202–5.

28. Bekolay T, Kolbeck C, Eliasmith C. Simultaneous unsupervised and supervised learning of cognitive functions in biologically plausible spiking neural networks. Cognitive Science Society; 2013. page 169–74.

29. Ringach DL. Spatial Structure and Symmetry of Simple-Cell Receptive Fields in Macaque Primary Visual Cortex. J. Neurophysiol. 2002;88(1):455–63.

30. Jones JP, Palmer LA. An Evaluation of the Two-Dimensional Gabor Filter Model of Simple Receptive Fields in Cat Striate Cortex. J. Neurophysiol. 1987;58(6):1233–58.

31. Camperi M, Wang X-J. A model of visuospatial short-term memory in prefrontal cortex: recurrent network and cellular bistability. J. Comput. Neurosci. 1998;5:383–405.

32. Miller P, Brody CD, Romo R, Wang XJ. A Recurrent Network Model of Somatosensory Parametric Working Memory in the Prefrontal Cortex. Cereb. Cortex. 2003;13(11):1208–18.

33. Palmeri TJ, Schall JD, Logan GD. Neurocognitive Modeling of Perceptual Decision Making. In: Busemeyer JR, Wang Z, Townsend JT, Eidels A, editors. Oxf. Handb. Comput. Math. Psychol. New York, NY: Oxford University Press; 2015. page 320–40.

34. Stewart TC, Bekolay T, Eliasmith C. Learning to Select Actions with Spiking Neurons in the Basal Ganglia. Front. Neurosci. 2012;6.

35. Fuster J. The prefrontal cortex. 5th ed. New York, NY: Elsevier; 2015.

36. Sreenivasan KK, Vytlacil J, D’Esposito M. Distributed and dynamic storage of working memory stimulus information in extrastriate cortex. J. Cogn. Neurosci. 2014;26(5):1141–53.

37. Stokes MG, Kusunoki M, Sigala N, Nili H, Gaffan D, Duncan J. Dynamic Coding for Cognitive Control in Prefrontal Cortex. Neuron. 2013;78(2):364–75.

38. Pasternak T, Greenlee MW. Working memory in primate sensory systems. Nat. Rev. Neurosci. 2005;6(2):97–107.

39. Rasmussen D, Eliasmith C. A spiking neural model applied to the study of human performance and cognitive decline on Raven’s Advanced Progressive Matrices. Intelligence. 2014;42:53–82.

40. Kajic I, Gosmann J, Stewart TC, Wennekers T, Eliasmith C. A Spiking Neuron Model of Word Associations for the Remote Associates Test. Front. Psychol. 2017;8:48.

41. Lewis-Peacock JA, Drysdale AT, Oberauer K, Postle BR. Neural evidence for a distinction between short-term memory and the focus of attention. J. Cogn. Neurosci. 2012;24(1):61–79.

42. Oberauer K. Design for a working memory. In: Ross BH, editor. Psychol. Learn. Motiv. Academic Press; 2009. page 45–100.

43. Cowan N. Attention and memory: An integrated framework. New York: Oxford University Press; 1995.

44. McElree B. Working memory and focal attention. J. Exp. Psychol. Learn. Mem. Cogn. 2001;27(3):817–35.

45. Anderson JR. How Can the Human Mind Occur in the Physical Universe? New York: Oxford University Press; 2007.

46. Nijboer M, Borst JP, Van Rijn H, Taatgen NA. Contrasting Single and Multi-Component Working-Memory Systems in Dual Tasking. Cognit. Psychol. 2016;86:1–26.

47. Borst JP, Taatgen NA, Stocco A, Van Rijn H. The Neural Correlates of Problem States: Testing fMRI Predictions of a Computational Model of Multitasking. PLoS ONE. 2010;5(9):e12966.

48. Borst JP, Taatgen NA, Van Rijn H. What Makes Interruptions Disruptive? A Process-Model Account of the Effects of the Problem State Bottleneck on Task Interruption and Resumption. Proc CHI. Seoul, Korea: ACM Press; 2015.

49. Larocque JJ, Lewis-Peacock JA, Postle BR. Multiple neural states of representation in short-term memory? It’s a matter of attention. Front. Hum. Neurosci. 2014;8:5.

50. Singh R, Eliasmith C. Higher-dimensional neurons explain the tuning and dynamics of working memory cells. J. Neurosci. 2006;26(14):3667–78.

51. Tripp BP, Eliasmith C. Population models of temporal differentiation. Neural Comput. 2010;22(3):621–59.

52. Eliasmith C, Anderson CH. Neural Engineering: Computation, Representation, and Dynamics in Neurobiological Systems. Cambridge, MA: The MIT Press; 2002.

53. Peirce J, Gray JR, Simpson S, MacAskill M, Höchenberger R, Sogo H, et al. PsychoPy2: Experiments in behavior made easy. Behav. Res. Methods. 2019;51(1):195–203.

